# Structural heterogeneity in biliverdin modulates spectral properties of Sandercyanin fluorescent protein

**DOI:** 10.1101/2021.04.02.438172

**Authors:** Swagatha Ghosh, Sayan Mondal, Keerti Yadav, Shantanu Aggarwal, Wayne F. Schaefer, Chandrabhas Narayana, S. Ramaswamy

**Author notes:** Authors contributed equally to this work (Swagatha Ghosh, Sayan Mondal). Department of Chemical Physics, School of Chemistry, Tel Aviv University, Tel Aviv 69978, Israel (Sayan Mondal).

## Abstract

Sandercyanin, a blue homo-tetrameric lipocalin protein purified from Canadian walleye (*Stizostedion vitreus*), is the first far-red fluorescent protein reported in vertebrates(1–3). Sandercyanin binds non-covalently to biliverdin IXα (BLA) and fluoresces at 675nm on excitation at 375nm and 635nm(1). Sandercyanin fluorescence can be harnessed for many *in vivo* applications when engineered into a stable monomeric form. Here, we report the spectral properties and crystal structures of engineered monomeric Sandercyanin-BLA complexes. Compared to wild-type protein, monomeric Sandercyanin (∼18kDa) binds BLA with similar affinities and show a broad red-shifted absorbance spectra but possess reduced quantum efficiency. Crystal structures reveal D-ring pyrrole of BLA rotated around the C14-C15 bond, which is stabilized by neighboring aromatic residues and increased water-mediated polar contacts in the BLA-binding pocket. A tetrameric Sandercyanin variant (Tyr-142-Ala) co-displaying red- and far-red absorbing states, and reduced fluorescence shows similar conformational changes in BLA binding pocket. Our results suggest that D-ring flexibility of BLA and its rearrangement reduces the fluorescence quantum-yield of monomeric Sandercyanin. Structures of monomeric Sandercyanin could be utilized as prototypes to generate bright BLA-inducible fluorescent proteins. Further, our study postulates a mechanism for modulating photo-states in BLA-bound lipocalins, known only in phytochromes till date.

**Significance Statement:** Sandercyanin is a tetrameric red fluorescent protein from a blue variant of walleye (*Stizostedion vitreum)* that binds to biliverdin IXα (BLA). Its biophysical properties and structures have been published earlier(1). A bright and stable monomeric Sandercyanin could be utilized as a fusion protein for fluorescence-based applications. Here we report the first structures and spectral properties of fluorescent monomeric Sandercyanin-BLA complexes and describe the molecular basis of modulated spectral properties due to rotated D-ring pyrrole around C14-C15 bond and re-shuffling of BLA-binding pocket. BLA-bound monomeric Sandercyanin could be engineered into brighter variants for *in-vivo* applications. Our study also reveals an unfamiliar mechanism in BLA-binding lipocalins that regulates red- and far-red absorbance states.

## Introduction

Sandercyanin is a blue colored protein found in the skin mucus of some North American walleye, *Stizostedion vitreum* (formerly *Sander vitreus*) (3). It is the smallest red-fluorescent protein (by monomer size) known in vertebrates that uses biliverdin IX-alpha (BLA) as a chromophore (1, 2). We have hypothesized the significance of Sandercyanin as a possible sunscreen to protect walleye, from increased exposure to harmful UV-radiation due to the ozone hole over the North Pole. Sandercyanin may also function in camouflaging walleye against their predators (1, 3). Sandercyanin exists as a homo-tetrameric protein and binds BLA by non-covalent interactions inside a lipocalin like beta-barrel fold. BLA-bound Sandercyanin absorbs at 375nm (UV) and 630nm (red) and shows far-red fluorescence at 675 nm on excitation with either wavelength. In the chromophore-binding pocket, BLA is stabilized by aromatic stacking interactions, hydrophobic interactions and H-bonding network from the surrounding amino acids residues(1). These interactions are naturally fine-tuned in Sandercyanin to give rise to a large spectral shift of 300nm to far-red fluorescence upon UV excitation at 375 nm(1).

Far-red fluorescence of Sandercyanin has potential to be employed for *in vivo* applications in biology and medicine(4–7). BLA-bound fluorescent proteins (FPs) such as bacteriophytochromes (BphPs) are used for near-infrared (NIR) imaging of deep-tissue due to their absorbance and emission spectral window, high signal to noise ratio, less scattering due to tissue transparency and less auto fluorescence from cells(4, 6). The abundance of BLA as a chromophore in mammalian cells makes BLA-binding fluorescent proteins, highly tractable to be engineered for next generation NIR FPs based applications(4). Oligomerization and low fluorescence quantum yield of Sandercyanin limit its ready-made utilization as a fluorescent protein marker(8). Hence, there remains a quest for engineering Sandercyanin to bright monomeric forms, and its further development for live cell imaging applications. Structure-based engineering and evolution of proteins have been successfully employed to tune FPs to obtain desirable properties(9–12).

Here we report engineered variants of tetrameric wild-type Sandercyanin into stable monomeric fluorescent forms using structured-guided site-directed mutagenesis. The three-dimensional structures of the variants, along with resonance Raman Spectroscopy and hybrid quantum mechanical/molecular mechanical (QM/MM) studies provide information on the correlation between the structure and observed spectroscopic properties.

## Results

### 1. Structure-based engineering yield stable BLA- bound monomeric fluorescent variants of Sandercyanin

The previously reported crystal structure of tetrameric Sandercyanin was used to identify residues at the dimeric interfaces that are critical for oligomerization (1), (Figure S1A). Single mutation of some of these residues generated stable monomeric-BLA complexes. Blue-green color of BLA-bound monomeric Sandercyanin variants were identified by size-exclusion chromatography at physiological conditions (Figure 1A). We selected two BLA- bound fluorescent monomeric mutants: Val-71-Glu (V71E) and Leu-135-Glu (L135E) (one from each dimer interface), for structural and functional analysis.

**Figure1.**
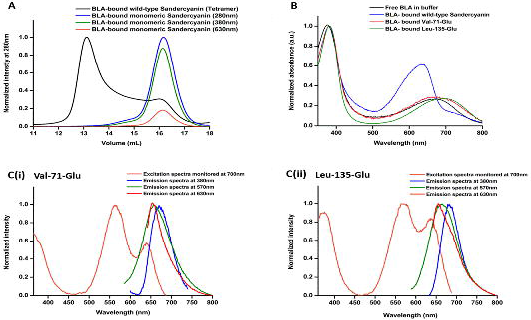
Characterization of monomeric Sandercyanin: (A) Size-exclusion chromatography profile of BLA-bound monomeric Sandercyanin (measured at different wavelengths), compared to wild-type tetrameric protein (black). (B) Overlapped absorbance spectra of BLA-bound monomeric variants (V71E and L135E), wild-type protein and free BLA. (C) Excitation and fluorescence spectra (in far-red region) of BLA-bound (i) V71E and (ii) L135E at different excitation wavelengths.

Absorbance spectra of BLA-bound V71E and L135E show maxima at 380nm and an attenuated red-shifted broad peak at far-red wavelengths (Figure 1B). Excitation spectra show splitting of the lower-energy band compared to wild-type Sandercyanin(1). Excitation of these monomeric variants at 380nm, 570nm and 630nm yield red fluorescence with maxima between 650 and 680nm for different variants (Figure 1C). No fluorescence is observed on excitation at far-red wavelength 700nm. Fluorescence quantum yield is diminished on monomer formation, however the binding efficiencies (K_d_) of BLA are similar to wild-type Sandercyanin (Figure S1B, Table S1). Since free BLA show enhanced red fluorescence in hydrophobic and aprotic solvents, we speculated that loss of oligomerization in these monomer variants possibly exposes BLA from Leu135- and Ala137-promoted hydrophobically locked state (at the dimeric interface) to aqueous/ protonated environment (Figure S1Ai) (1, 13). Addition of a detergent (Triton X-100) increased the peak fluorescence of monomeric variants by 10-20%, suggesting a crucial role of hydrophobicity in maintaining quantum yield of BLA-bound monomeric Sandercyanin (Figure S1C).

### 2. Crystal structures of monomeric variants elucidate the molecular basis of altered spectral properties of Sandercyanin

We obtained BLA- bound crystals of V71E and L135E at 277K and dark conditions to avoid any photo-damage to the crystals (Figure S2A). The structures of V71E and L135E were determined at 2.5Å and 2.75Å respectively, using wild-type protein structure as model for molecular replacement. Data collection and refinement statistics are summarized in Table S2. Both monomeric variants show a lipocalin fold forming a beta-barrel, similar to a monomeric subunit of wild type protein, with BLA bound at the center of the barrel (Figure S2B). Loss of oligomerization creates a vacant space in the monomer near the open/ wider end of the barrel, exposing BLA at the site of dimerization. N-terminal amino acids and protein backbone of loop residues (Leu50-Glu58, Arg80-Ile86, His108-Val114 and Ile 133-His139) are perturbed and some of them appear to fold inwards in the monomer (Figure S2B).

In both variants, D-ring pyrrole of BLA rotated around the C14-C15 single bond with a torsion angle of 115° (counter-clockwise) compared to its native ZZZssa configuration in wild-type Sandercyanin (Figures 2A, 2B). While V71E purely maintains a ZZZsss conformer, we found the possibility that both ZZZsss and ZZEsss conformations (resulting from intrinsic rotation and isomerization of the D-ring around C14-C15 and C15=C16) fit the electron density in L135E with similar B-factors of both conformations. Aromatic residues Phe-55, Phe-106, His-108 and Tyr-142 undergo concerted rearrangement of side chains to maintain stacking with the rotated D-ring pyrrole and confining it to a sterically hindered position (Figure 2C). The two monomeric variants have distinct changes in the conformation of the re-oriented aromatic residues. Non-aromatic amino acids near BLA show subtle structural changes. The C-ring and its propionic chain get displaced with the overall chromophore moving slightly upwards from wild-type BLA (Figure 2D). There are more water molecules near BLA forming polar contacts with the rotated chromophore. Water molecules, substitute the reoriented side-chains of Phe-55 and Tyr-142. A water molecule is stabilized within the BLA pyrroles and the carbonyl group of the rotated D-ring forms water-mediated H-bonds with Asp-47, Gln-56, Asn-53 and Ser-144 (Figure 2E). Newly adopted conformations of BLA, stabilized by neighboring aromatic residues and more polar contacts in the chromophore-binding pocket causes far-red absorbance (SFr) in monomeric Sandercyanin and possibly lowers quantum yield of fluorescence.

**Figure 2.**
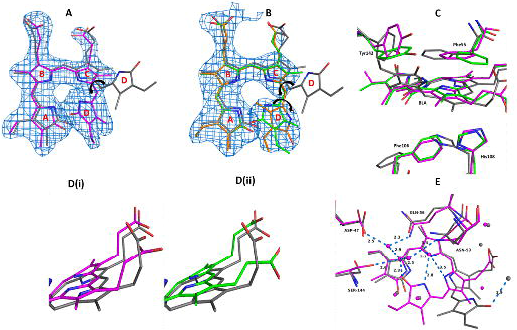
Structural insights of BLA-bound monomeric Sandercyanin: 2Fo-Fc maps (contoured at 1σ) of BLA (blue) in the binding pocket of (A) V71E and (B) L135E, showing rotated D-ring pyrrole around C14-C15 bond forming ZZZsss (magenta, green) or ZZEsss (orange) compared to and wild-type protein ZZZssa isoform (grey). (C) Rearrangement of aromatic residue with the binding pocket stabilizes BLA in its new conformation. (D) Perturbation in propionic group of C-ring and upward displacement of BLA and (E) increased number of water molecules (non-bonded spheres in magenta and grey, respectively) in BLA-binding pocket of V71E (magenta) making more polar contacts and water-mediated H-bonds with BLA compared to wild-type protein (grey). Similar observation was made in L135E (SI appendix, Figure S2C).

### 3. Tetrameric variant (Tyr-142-Ala) reveals structural-heterogeneity in the BLA-binding pocket, similar to monomeric Sandercyanin

To decipher the role of conformational flexibility of BLA on spectral properties of Sandercyanin, we generated alanine (ala)-variants of aromatic residues (Phe-106, His-108 and Tyr-142) in wild-type protein. All variants form tetramer on binding to BLA, but exhibit differences in their absorbance spectra compared to wild-type protein (Figure 3A). His-108-Ala show increased intensity of 720nm band, without shift in peak at 630nm, while Phe-106-Ala show minimal deviation in absorption spectra compared to wild-type Sandercyanin. A shift in maxima from 630nm to 660nm is observed in Tyr-142-Ala (Y142A), with concomitant appearance of the 720 nm far-red band with similar intensity. On excitation with 380nm and 630nm, Y142A show fluorescence with maxima at 685nm and fluorescence quantum yield equivalent to monomeric V71E variant (Figure S3A, Table S1).

**Figure 3.**
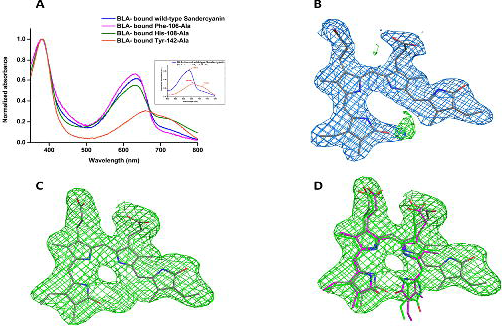
Spectral properties and structure of BLA-binding pocket of Y142A: (A) Tetrameric variants of Sandercyanin with alanine mutation of aromatic residues show presence of far-red state in the absorbance spectra. (B) 2Fo-Fc map (blue, contoured at 1σ) of BLA in ZZZssa conformation (grey) in Y142A showing residual Fo-Fc density (green, contoured at 3σ) at the center of BLA. (C) Unbiased Buster map (green, contoured at 3σ) of ligand-free Y142A showing presence of extra density around BLA and (D) accommodating BLA conformations from V71E (magenta) and L135E (green) structures.

Crystal structure of Y142A variant was determined to 2.65 Å with BLA in the (native) ZZZssa configuration displayed residual positive Fo-Fc density between the A- and D-ring pyrrole, indicating partial occupancy of the chromophore at that position (Figure 3B). Using a ligand-free refined structure of BLA- bound Y142A, we calculated an unbiased Buster-map that revealed additional Fo-Fc density of BLA in the binding pocket (Figure 3C) (14, 15). The new Fo-Fc density accommodates the rotated ZZZsss (or ZZEsss) conformations of BLA in monomeric variants of Sandercyanin (Figure 3D). Assigning occupancy of 25% to an alternate conformation (ZZZsss) of BLA gives acceptable fit with similar B-factors. Refinement with partially occupied ZZZsss conformation generates density for water molecule (water-93), overlapping with water-68 in the monomeric variants forming water-mediated H-bond with the rotated pyrrole (Figure S3B). Further, mutated Tyr-142 accommodates a water molecule (water-86) in its position, similar to water-67 in V71E structure (Figure S3B). Our observations suggest that BLA has a more flexible D-ring in Y142A and show increased polar contacts in the protein matrix compared to wild-type Sandercyanin, that in-turn, reduces the fluorescence quantum yield of Sandercyanin. Further, to elucidate role of conformational heterogeneity in producing co-existing red (Sr; 660nm) and far-red (SFr; 720nm) photo-states of Y142A, we employ solution state Raman spectroscopy and theoretical modelling (*vide infra*).

### 4. Resonance Raman and computational studies decipher conformational plasticity of BLA in Sandercyanin

Solution state resonance Raman (RR) spectroscopy in two electronic states (Figure 4A, top, inset), and molecular dynamics (MD) and QM/MM computation allowed us to probe conformations of BLA in Sandercyanin(16, 17). The RR spectra of free- and bound BLA to wild-type protein at 405 nm (Figure 4A, 4B top) and at 532 nm (Figure 4A, 4B bottom) reveal that binding to Sandercyanin leads to spectral narrowing of majority of vibrational bands of BLA, and only affect positions in the high frequency region that represent primarily the resonance enhanced in-plane vibrations coupled to D-ring. (Figure S4, and Table S3 for normal mode assignments of all bands). The loss of RR intensity in these bands imply sudden decrease in polarizability of these modes – probably due to disruption of pi-electron conjugation, and suggesting possible rotation of D-ring of BLA and stabilization of a specific BLA conformation inside Sandercyanin(18).

**Figure 4.**
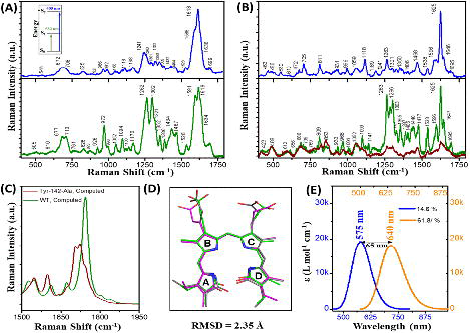
Resonance Raman spectra and molecular simulation probe D-ring rotation of BLA in Y142A. RR spectra of (A) free BLA, and (B) wild type Sandercyanin with 405 nm (top) and 532 nm (in green, bottom) excitation, and of Y142A (in brown, bottom) with 532 nm excitation. (C) QM/MM computed normal Raman spectrum of WT (green) and Y142A (brown) Sandercyanin. (D) Superimposed BLA conformation obtained from MD (magenta), MC-SD (green) simulation, and crystal structure (grey). (E) TDDFT//QM/MM computed absorption spectrum of most (orange) and second most (blue) populated structure obtained from MD (Figure S6). Two spectra in panel B (bottom) are normalized at 770 cm^−1^ (marked by arrow). Schematics of the energy diagram of two electronic states of BLA probed here are shown as inset of panel A.

In contrast to wild-type protein, BLA bound to Y142A shows loss of fine structures of several RR bands in 532 nm excited spectra, specifically of four bands at 1695, 1647, 1625 and 1596 cm^−1^ associated with D-ring stretching vibrations (Figure 4B, bottom, in brown and see Figure S5). Significant broadening of these RR bands strongly suggests occurrence of more than one BLA conformers when bound to Y142A in solution state. Our MD simulation reveals rotation of BLA D-ring around C14-C15 bond, and stabilization of ZZZsss conformation upon mutating Tyr-142 to Ala (Figure S6, See Supplementary information, SI, for details of simulations, and Videos S1, S2). To probe conformational space of BLA owing to its intrinsic flexibility inside Y142A, hybrid Monte Carlo-stochastic dynamics (MC-SD) sampling of all torsional coordinates around each single bond of BLA was performed (Figure S7A). Majority (79%) of the conformers yields the same core structure as obtained from MD simulation, and also agreeing with the crystal structure with a RMSD of 2.35 Å (Figure 4C).

Large reduction of RR intensity in 1200-1750 cm^−1^ region not only indicates mere loss of pi-electron conjugation between adjacent covalently linked rings, but also disruption of non-covalent pi-pi interactions with aromatic residues stabilizing BLA. We support our claim through computing Raman activity of Y142A, and wild-type protein. Computed spectrum (on the wild-type, and the structure described in Figure S6E) shows loss of intensity of the vibrational modes that are associated with in-plane D-ring vibrations. (Figure 4D, and Figure S5B) QM/MM computation on representative structure from five clusters (Figure S6A-E) of Y142A shows that heterogeneous structural ensembles resulting in two similarly intense (oscillator strength = ∼0.25) absorption bands at ∼578 nm and 640 nm, separated by 62 nm (Figure 4E and Table S4). This is in very good agreement with experimental observation of ∼60 nm difference between Sr state at 660nm and SFr state at 720 nm in Y142A (Figure 3A). In-line with experiment, ^1^ππ* state of wild-type protein (on PDB:5EZ2 model) is predicted at 568 nm with a much higher (compared to that of Y142A models) oscillator strength of 0.45 corresponding to the experimental band at 630nm. Thus, our models show that conformational heterogeneity not only induces the red shift of the ^1^ππ* transition to produce SFr state in Y142A, but also leads to observed loss of absorption cross-section.

## Discussion

In this report we present the structures of BLA- bound monomeric Sandercyanin fluorescent proteins and illustrate the molecular basis of altered spectral properties by extending our study to a tetrameric variant of Sandercyanin. Structure-based directed mutagenesis have been employed successfully for engineering of fluorescent protein for their utilization in *in vivo* imaging applications(9, 10, 12, 19, 20). Using previously reported structures (PDB code: 5EZ2 and 5FZ6), we generated site-specific modification of amino-acids involved in oligomerization of wild-type Sandercyanin. Similar binding affinities of monomer and wild-type protein suggest ligation of BLA to Sandercyanin is independent of the oligomeric state of the protein. BLA- bound monomeric Sandercyanin with molecular weight of ∼18KDa are till date the smallest far-red fluorescent protein known in eukaryotes(1, 21).

Spectral and structural studies show two major factors responsible for reduced fluorescence quantum yield of monomeric Sandercyanin. Firstly, exposure to protic environment in the BLA-binding pocket (due to increased water molecules) play significant role in quenching fluorescence of chromophore (1, 13, 22). We have reported that increasing hydrophobicity around free BLA and deuteration of wild-type Sandercyanin increases the fluorescence intensity, possibly due to reduced nonradiative excited-state proton transfer (ESPT) processes in Sandercyanin(1). Secondly, rotation of D-ring of BLA to adopt multiple ground-state conformations (ZZZsss and ZZEsss) could affect quantum yield of Sandercyanin. Fluorescent proteins (FPs) developed from BLA-binding bacterial phytochromes (BPhs) recognized conformational flexibility of BLA as an important factor for reduced fluorescence quantum yield(23–25). Due to several conformations of BLA observed in monomeric protein (also evident by its broad absorbance spectra) and Y142A variants, it is possible that only a fraction of BLA conformers (mainly the ZZZssa isoform) undergo radiative transition to show fluorescence.

Structural changes such as rearrangement of aromatic amino acids, dislocation of C-ring and upward-sliding of BLA could also influence the optical properties of BLA in Sandercyanin. It is evident from crystal structures that Phe-106 and Tyr-142 play crucial role in maintaining stacking interactions with the D-ring, while Phe-55 (positioned over BLA in wild-type protein) and His-108 stabilizes the C-ring. Propionic side-chains of B- and C-ring pyrroles of BLA have been shown to affect photoconversion in bacteriophytochromes from *Agrobacterium fabrum*(26). Our computational studies also suggest that dynamic interaction between His-108 and C-ring of BLA in Y142A play a crucial role in stabilizing the Sr and SFr states (Figure S6A-E). Hence, we infer that perturbations induced by aromatic side chains of binding site residues to C- and D-ring regulates spectral properties of Sandercyanin.

Ground state structural heterogeneity due to D-ring rotation around the C14-C15 single bond promotes transition from Sr to SFr photo-states in Sandercyanin. This mechanism is seldom known in lipocalins and non-phytochromes. Sandercyanin and bacterial phytochromes (BPhs) bind to the ZZZssa conformation of BLA in the dark (or resting) state of the protein, yielding red-absorption state ∼630-645nm (1, 27, 28). BPhs undergo photo-isomerization of the C15=C16 double bond of BLA from ZZZssa to ZZEssa conformations, giving rise to red (Pr)- and far red (Pfr)-respectively (27–30). Recent reports showed a twisted D-ring rotation coupled with partial detachment of BLA from the protein as a primary photo-response in a BPh from *Deinococcus radiodurans* observed at time-scales of 1picoseconds (ps)(31). Further, two Pr ground states of BLA have been reported in a cyanobacterial phytochrome Cph1 as a result of altered charge distribution and hydrogen bond network near ZZZssa state(32). Light-induced photoconversion is not yet established in Sandercyanin. However, partial occupancy of two BLA conformers seen in crystal structure and in molecular simulations of Y142A explain Sr and SFr photo-states in Sandercyanin.

While crystal structure shows BLA in a ZZZsss sate, our simulations reveal further structural heterogeneity responsible for two observed electronic transitions in Y142A. The major population of have a stabilizing stacking interaction between C-ring and His-108 responsible for the SFr state. (Figure S6E). Reorganization of residues, specifically partial de-stacking of His-108 and Phe-55 stabilizes the other ensemble leading to Sr photo-state. (Figures S6A-D, and S6F). Interestingly, the most stable conformation of BLA in tetrameric Y142A overlaps with that in V71E monomeric variant, and is found to be stabilized by residues in the binding pocket, as seen in crystal structures (Figures 2A, 2C, 2E, and S7C). Thus, our hybrid approach provides critical insight that ground-state conformational heterogeneity in Y142A originates at two levels; (1) intrinsic flipping of D-ring due to reduced steric hindrance, and (2) reorganization of the binding pocket residues to stabilize a particular BLA conformation. Interestingly, A-ring pyrrole of BLA, unlike BPhs, associate non-covalently in Sandercyanin and possess similar conformational freedom as D-ring, but remains unperturbed in all variants studied here. Thus, our results demonstrate a dominant structural characteristic of D-ring pyrrole in driving chemistry of BLA- or linear-tetrapyrroles binding proteins across diverse species(33, 34).

In conclusion, our study demonstrated chemo-spatial characteristics of the chromophore-binding pocket of monomeric Sandercyanin-BLA complexes, and is crucial for development of bright far-red fluorescent proteins for cellular studies(10, 19, 35–37). Sandercyanin, is unique as a fluorescent protein and challenging to modify, and with only a few engineered lipocalins reported till now(38, 39). Given our success of engineering monomer variants of Sandercyanin without losing the fluorescence, we propose that a bright monomeric version could be generated with increasing the hydrophobicity and attaining a sterically-locked ZZZssa conformer of BLA in Sandercyanin(7, 25). As Sandercyanin is a eukaryotic protein, there are innumerable opportunities for utilizing Sandercyanin for cellular-applications, where BLA is produced intrinsically as a metabolite(4, 6, 40).

## 3.6 Methods and materials

The details cloning, protein purification, biophysical assays, crystallization and structure determination, resonance Raman spectroscopy, and molecular simulations are described in SI.

## 3.5 Acknowledgments

The preparatory work for this project, spectroscopy and crystallographic data analysis was carried out at Institute for Stem Cell Science and Regenerative Medicine (inStem, Bangalore, India). Crystallography data collection was performed at BM14, ID23 and ID30A-1 beamlines at ESRF (Grenoble, France). We thank all members of our laboratory at inStem (India) for their support during the course of this research. Further, we thank beamline staff at ESRF for help in data collection. Support for this work was provided from Department of Biotechnology India (Grant BT/PR5801/INF/22/156/2012) and WISYS Technology Foundation, Inc. (Madison, WI, USA). C.N. would like to acknowledge LSRET-JNC (BT/INF22/SP27679/2018) intramural funds for financial assistance.

## Author contributions

S.G. and R.S. planned the research. S.G. designed and characterized the clones for detailed studies. S.G. and K.Y. purified protein, performed UV-VIS spectroscopy, protein crystallization and analyzed crystallography data. S.M. and S.A. carried out RR data-collection; S.M. performed detailed data analysis, interpretation of RR data and all theoretical studies. R.S., C.N., W.F.S and S.G supervised the project. S.G. and S.M. wrote the manuscripts and received inputs from all authors.

## Conflict of interest

The authors declare that they have no conflict of interest.

## Protein structure deposition

The atomic coordinates and structure factors have been deposited PDB IDs: 7O2Y (V71E), 7O3K (L135E) and 7O32 (Y142A) in the Protein Data Bank (www.pdb.org).

## Notes

### Competing Interest Statement

The authors have declared no competing interest.

