## Supplementary material for "Structural heterogeneity in biliverdin modulates spectral properties of Sandercyanin fluorescent protein": Sandercyanin_SI_Ghosh_et_al_020421.docx

**SUPPLEMENTARY INFORMATION (SI)**

### 1. Site-directed mutagenesis, protein expression and purification of Sandercyanin variants.

To design monomer mutants of Sandercyanin, we identified amino acids involved in oligomerization of the wild-type protein using high resolution X-ray crystal structures (1) (Figure S1A). We have previously reported about BLA- induced oligomerization of Sandercyanin from monomeric apo protein in solution to a tetrameric BLA- bound form. Leu-135, Ala-137 and Ser-138 emerging in a long loop from one monomer interact with BLA of neighboring monomer which possibly results in BLA- induced tetramerization of wild-type protein. Amino acid residues: Ser-68, Pro-69, Val-71, Ile-94, Glu-96, Asp-97, Pro-98, Glu-109 and Asn-110 are involved purely in protein- protein interaction at the other dimer interface. Selected residues were mutated to change the physical and chemical property of the amino acids and the nature of interaction holding the interface. Further, using structures of monomeric variants Val-71-Glu (V71E) and Leu-135-Glu (L135E), we targeted aromatic amino acids at the ligand- binding pocket in wild-type tetrameric Sandercyanin to induce conformational changes in BLA. The gene for wild type Sandercyanin cloned into pET21a (+) was used as a template for generating site- specific mutations. We performed whole- vector polymerase chain reaction (PCR) of the template gene using high- fidelity Phusion DNA polymerase (New England Biolabs) (Ref). Verification of correct mutation was confirmed by sequencing of plasmid purified from DH5α *E. coli* cells and subsequent alignments of mutated sequences with the wild-type protein using Clustal-W(2). Expression and purification of Sandercyanin variants were performed in using similar techniques as wild-type protein, as described previously(1). For monomeric and tetrameric variants, blue-colored BLA-bound fractions corresponding to size of ~18kDa and ~75kDa, respectively, were collected and used for spectroscopy and crystallographic studies. Apo-protein of each variant were purified using the same method without BLA.

### 2. Absorbance and fluorescence spectroscopy.

All experiments were performed with freshly purified protein samples (apo and holo forms) at pH 7.5 and room temperature. UV-Visible absorbance spectra of Sandercyanin variants were recorded with ultraspec 2100 pro spectrophotometer from Amersham Biosciences. Fluorescence and binding studies of Sandercyanin variants were monitored on Horiba Jobin Yvon Fluoromax-4 fluorimeter as described in previous report(1). Data analysis and plotting of spectra was performed using OriginPro8 software.

### 3. Crystallization, data collection and structural analysis.

Purified Sandercyanin variants are mixed with excess BLA before crystallization. Hanging drops were set-up at 4^º^C using mosquito high- throughput crystallization system from TTP life sciences and crystals were grown by vapor diffusion. BLA- bound protein crystals obtained in various conditions and different colors (Figure S2A) were flash-cooled in liquid nitrogen after soaking in 10% ethylene glycol as cryo-protectant. Crystallography datasets of monomeric Sandercyanin variants and Tyr-142-Ala (Y142A) were collected at ESRF (Grenoble). All images were indexed, integrated and scaled using HKL2000 (3) or d*TREK (4) or MOSFLM (5) and structure was determined using molecular replacement with Phaser (6) using wild-type Sandercyanin structure as starting model All structures were refined with CCP4i (7) and PHENIX (8). Presence of BLA in the protein was indicated by F_o_-F_c_ density in the BLA- binding pocket. Model building was done with Coot (9) and all structural illustrations were generated using PyMOL (10). All parameters of data collection and refinement statistics are summarized in Table S2.

### 4. Resonance Raman spectroscopy

Resonance Raman (RR) spectra of wild-type Sandercyanin and Y142A variant were recorded using LabRAM HR Evolution confocal Raman microscope (HORIBA Jobin Yvon S.A.S, France) equipped with a spectrograph of 800mm focal length and a Peltier-cooled CCD to register the spectra. A holographic diffraction grating with 1800 gr/mm, and a 300μm spectrograph entrance slit was used for high spectral resolution. All protein samples were prepared in phosphate buffer at pH 7.4 and of 250uM in concentration for RR measurements. Samples were loaded in EPR quality capillary made of suprasil quartz of outer diameter 3mm (Wilmad-LabGlass, NJ, USA) and RR spectra are recorded in 180º backscattering geometry. Samples were resonantly excited with 405nm or 532nm solid state laser operated in CW mode through an 50x objective. The laser power on sample was 4-5 mW for 405 nm, and 8-9 mW for 532 nm laser. Required laser power for good signal to noise ratio was optimized without photo damage of sample. With 405 nm excitation, each spectrum was recorded with 90 s exposure and a total 270 s accumulation. In case of 532 nm excitation, an exposure of 20 s along with a total 100 s accumulation was used to acquire a spectrum. Final spectrum was average of 3 to 5 spectra each recorded from a fresh sample. Recorded RR spectra are analyzed using LabSpec 5 and Origin 7.

### 5. Model generation, Molecular Dynamics, hybrid Monte Carlo, and quantum mechanical calculations

**Model for wild-type Sandercyanin**

The complete protocol of all used theoretical methods is described as flowchart in Fig. S8. Experimentally determined structure of recombinant wild-type Sandercyanin (PDB: 5EZ2) was used for the molecular dynamics (MD) and other quantum chemical computation.(11) Coordinates of one monomer chain was extracted, and all computations are performed on this monomer corresponding the wild-type Sandercyanin. The preparation of all Sandercyanin models were carried out with Protein Preparation Wizard of Schrodinger Suite of programs (Schrödinger LLC, NY, USA), and all molecular dynamics (MD) simulations were performed using the Desmond program implemented within Schrodinger Suite.(12) Quantum chemical computations were performed with Gaussian 09 suite of programs. (13) Charge on all residues are set using ProPka at pH 7.4,(14) and heavy atoms are then energy minimized with OPLS3 force field, and was later solvated in orthorhombic, periodic boxes (10 x 10 x 10 Å^3^) with explicit TIP4P water model.(15) Then the solvated Sandercyanin was charge neutralized by addition of appropriate number of Na^+^ or Cl^-^ ions by replacing randomly chosen water molecule, which are at least 8 Å away from the bound BLA and the residues that are interacting with the BLA.

**Model for Y142A Sandercyanin**

The starting structure for Y142A was created by mutating Tyr-142 to Ala on a monomer of wild-type Sandercyanin (PDB: 5EZ2) in Protein Preparation Wizard of Schrodinger Suite, and keeping everything intact. Evolution of this structure was followed with NPT MD as described below.

**Molecular Dynamics**

The solvated monomer of Y142A mutant were relaxed and equilibrated using default Desmond minimization and equilibration procedures before MD production run. In short, first, Brownian dynamics at 10K was carried out in an NVT ensemble for 100 ps, following another NVT simulation at 10K using Berendsen thermostat(16) relaxation constant of 100 fs for total 12 ps. Following this step, the system was simulated in the NPT ensemble at 10K temperature with 100 fs temperature relaxation constant, and at 1 atm pressure with a 50 ps pressure relaxation constant. NPT simulation was then performed at 300K temperature, and 1 atm pressure using for 12 ps. For all four mentioned stages, non-hydrogen heavy atoms of proteins were restrained with a 50 kcal/mol force constant. At the final stage, the system is simulated in NPT ensemble for 24 ps without any restraints at 300K temperature and 1 atm pressure using 100 fs temperature relaxation constant, and 2 ps pressure relaxation constant respectively. Berendsen thermostat and Berendsen barostat were used for all these five steps of equilibration protocol.

This relaxed and equilibrated system is followed in an NPT ensemble for production run at 300K temperature for 500 ns using the Martyna-Tobias-Klein (MTK) barostat (17) with a 2 ps relaxation time constant, isotropically coupled to the Noose-Hover thermostat(18) having a 1 ps relaxation time constant. MD simulations are carried out with periodic boundary condition. During simulation, long-range electrostatic energy and forces were treated with particle-mesh-based Ewald (PME) technique using a columbic cutoff radius of 9 Å. Corrections to particle positions and momenta were obtained using M-SHAKE algorithm.(19) A RESPA-based integration scheme is employed with a time step of 2 fs for computing bonded interactions, 6 fs for long-range electrostatic forces, and 2 fs for short-range non-bonded forces. The MD simulations were run on a custom workstation using an Nvidia Geforce GTX 1070 graphic processor. All analysis of MD trajectories were performed with python scripts within Schrödinger Python API.

In this MD trajectory, we observe intrinsic flipping of the D-ring in Y142A binding pocket after 480 ns, and remain in this flipped state for remaining of the simulation time (see Video S1, S2 showing this ring flipping from the trajectory). We choose one structure with flipped D-ring, (details described in Fig. S8) and performed an independent MD simulation for 50ns in an NPT ensemble with 20ps time step with all parameters as described in previous paragraph. This trajectory is clustered in five groups based on RMSD of all heavy atoms of BLA, and residues within 4 Å of BLA. From each cluster, one structure with the smallest RMSD from the average structure of that cluster is selected as representative of that cluster, described in Fig. S6. All QM/MM computations of Raman spectrum and photo absorption spectrum are performed on these representative structures.

**Monte Carlo/Stochastic Dynamics (MC-SD) Conformational Searching**

Possible BLA conformations inside the binding pocket of Y142A mutant of Sandercyanin are sampled with a constant temperature mixed Monte Carlo-Stochastic Dynamics (MC-SD) simulation.(20) D-ring rotation in BLA can occur via C14-C15 single bond between ring C and D. Thus, a rapid search of the configurational space was performed through torsional scan around this bond by MC-SD method. In this method, first, stochastic dynamics (SD) is used to generate a trajectory at 300K, then a large step was sampled by a standard MC method for the torsional degrees of freedom. This step is accepted or rejected based on metropolis criteria. In this hybrid method, while SD samples the small amplitude collective motions near the potential energy minima, standard MC samples the large amplitude motions involving occasional crossing along the torsion barrier. By using the alternating SD, followed by a MC step, MC-SD explores the conformational space along all the torsional degrees of freedom in BLA very fast.	Four Monte Carlo trials has been performed for each stochastic dynamic time step. Two torsions chosen randomly from the list of all specified ones are varied in each MC step. The leap-frog variant of velocity Verlet integrator was used with a timestep of 1.5 fs at 300 deg K in stochastic dynamics. No constraint was applied to any bonds during the dynamics. The system is equilibrated for 5 ps before a simulation of 200 ps.

**QM/MM computations**

We use multi-layer ONIOM (Our Own N-layered Integrated molecular Orbital and molecular Mechanics) based hybrid QM/MM method to compute Raman and photo absorption spectrum of wild-type and Y142A Sandercyanin.(21, 22) All QM/MM computations are performed as two layer ONIOM (QM-high : MM-Low) with Gaussian 09 suite of programs.(13) Whole BLA, and side chains of residues Phe-55, Gln-56, Asn-77, Lys-87, His-108, Ala-142 residues are placed in the QM-high region, and treated with M06-2X functional,(23) and with 6-31g(d) basis set, while remaining of the proteins are included in the MM-low layer, and treated with amber force field. (24) M06-2X functional was previously shown to describe non-covalent interactions including hydrogen-bonding ones in protein system.(25–27) We used RESP v2.2 in R.E.D. tools package to compute the Amber charges for the atoms of the BLA chromophore.(28) Force field parameters for BLA were obtained using the Parmchk2 module of AmberTools based on the generalized Amber force field (GAFF).(24) For all ONIOM calculations, the high-QM layer has been separated from low-MM layer with hydrogen-capped link atoms placed at C-N bond interface between two amino acid residues. (see Fig. S8, inset) On the ONIOM(M06-2X/6-31g(d)//Amber) optimized ground state structure, harmonic vibrational analysis and Raman activities are computed for the high layer within the two layer ONIOM framework.(21) On this structure, vertical S_1_ ← S_0_ transition energy is computed with ONIOM(TD-DFT:AMBER) with 6-31+G(d) gaussian basis set on QM-high layer.(29–31) We used the hybrid exchange–correlation functional CAM-B3LYP (32) because it has shown to perform extremely well for excited states of biological chromophore, such as retinal (33), and also for predicting protein induced red shift in the absorption spectrum of GFP. (34) Electrostatic interaction between the BLA and the residues residing in QM-high layer, and the surrounding hydrophobic protein environment treated at MM-low level is taken into account via Electronic embedding that uses the partial MM charges of the low layer into the QM Hamiltonian of high layer.(32) The TD-ONIOM computed oscillator strengths are deconvoluted with gaussian line shape with FWHM of 3000 cm^-1^ to construct the photo absorption spectrum, and the computed Raman spectra are generated with Lorentzian line shape of FWHM of 25 cm^-1^.

**Normal Modes of Biliverdin**

All observed RR bands of free BLA, wild-type and Y142A Sandercyanin, and their assignments in terms of BLA normal modes are described in detail in SI Appendix, Table S4. This complete band assignments of BLA are done with our vibrational analysis at RB3LYP/6-31+G(2d,p)//PCM level of DFT, and previously published results of normal mode analysis of biliverdin derivatives, (35–37) and of phytochromes.(38, 39)

**SUPPLEMENTARY FIGURES AND TABLES**

**Figure S1: Structure-based engineering of Sandercyanin to monomeric variants and their properties**

(A) Crystal structure of tetrameric wild-type Sandercyanin showing (i) BLA of one monomeric subunit interacting with amino acids from neighboring subunit and (ii) second dimeric interface made purely by protein- protein interaction. (B) Binding curve of BLA to Sandercyanin variants: V71E, L135E and Y142A, determined using fluorescence measured at 680nm on excitation with 380nm (or 630nm). (C) Titration of BLA- bound monomeric Sandercyanin variants with a detergent (TritonX-100) showing increasing of fluorescence by 10-20% on addition of a hydrophobic medium around monomeric protein.

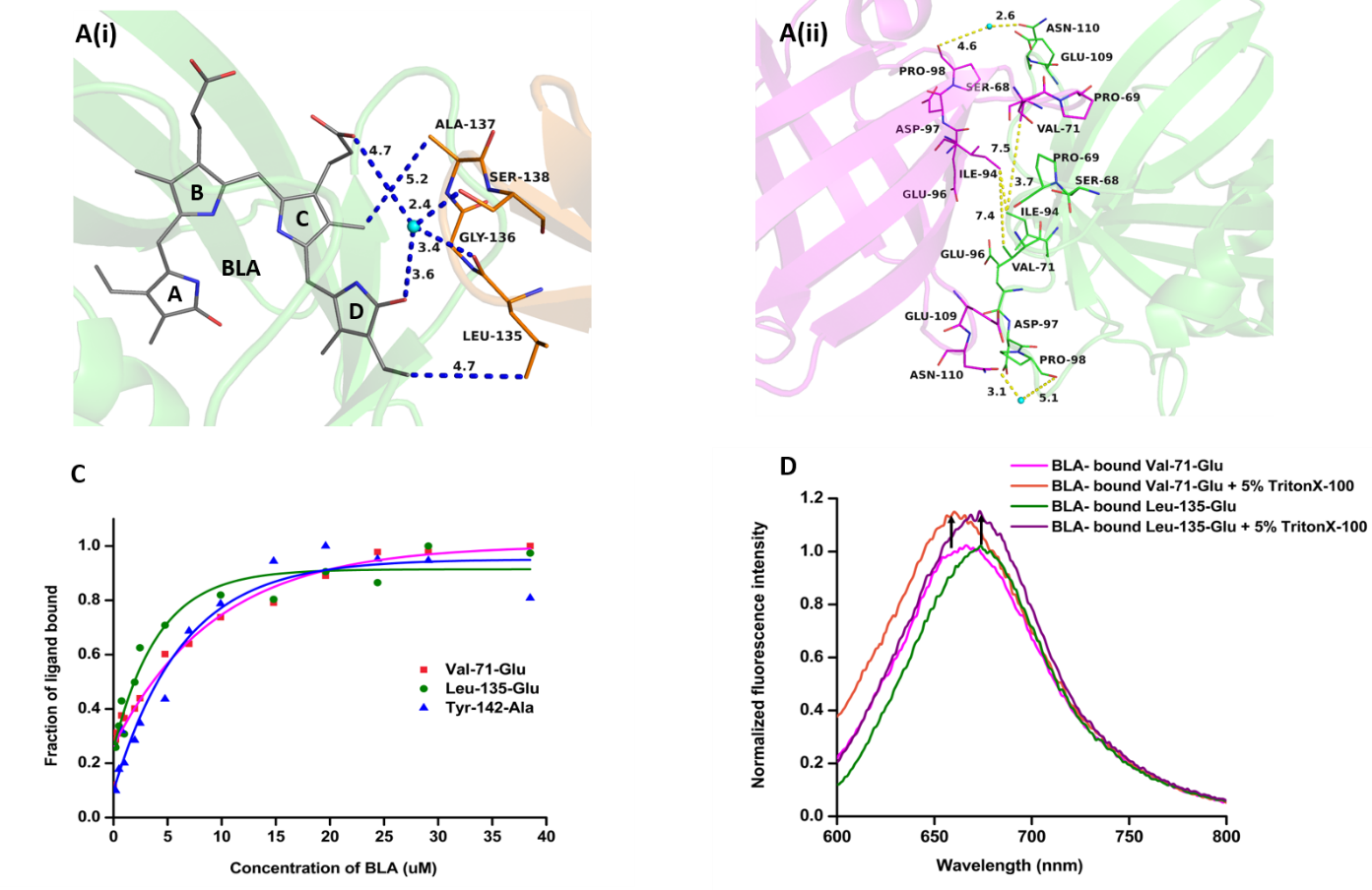

**Table S1: Spectral properties and binding efficiencies of monomeric variants of Sandercyanin**

| **Mutant name** | **Molecular weight** | **Ex/ Em1** | **Ex/ Em1** | **Kd** | **Quantum yield** |
| --- | --- | --- | --- | --- | --- |
| Wild-type Sandercyanin | 74.5 kDa | 375/675 | 630/ 675 | 6μM | 0.016 |
| V71E | 18.6 kDa | 380/670 | 600/660 | 5.1μM | 0.003 |
| L135E | 18.6 kDa | 380/682 | 600/676 | 2.7μM | 0.0025 |
| Y142A | 74.5 kDa | 380/685 | 580/653 | 11.1μM | 0.004 |

**Figure S2. Crystal structure of monomeric variants elucidate the molecular basis of far- red absorbance in Sandercyanin**

(A) Crystals and crystallization conditions of Sandercyanin variants in complex with BLA. (B) Overlapped structures of BLA-bound monomeric variants: V71E (magenta) and L135E (green) of Sandercyanin with monomeric subunit of wild-type protein (grey) showing a lipocalin fold and perturbation in the loop residues (yellow arrows) as a result of monomer formation. (C) BLA- binding pocket of L135E (green) showing increased number of polar contacts with the chromophore compared to wild- type protein (grey).

(A)

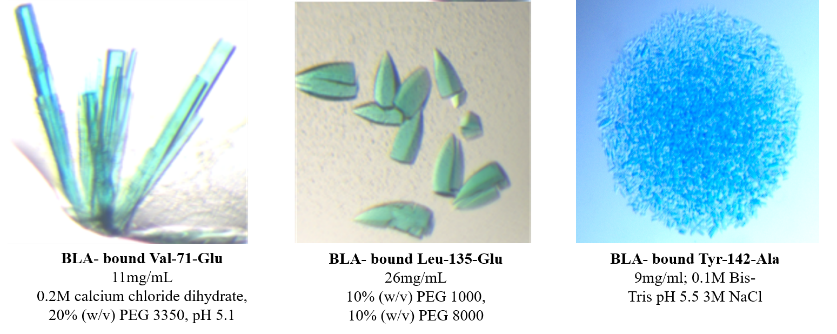

(B)

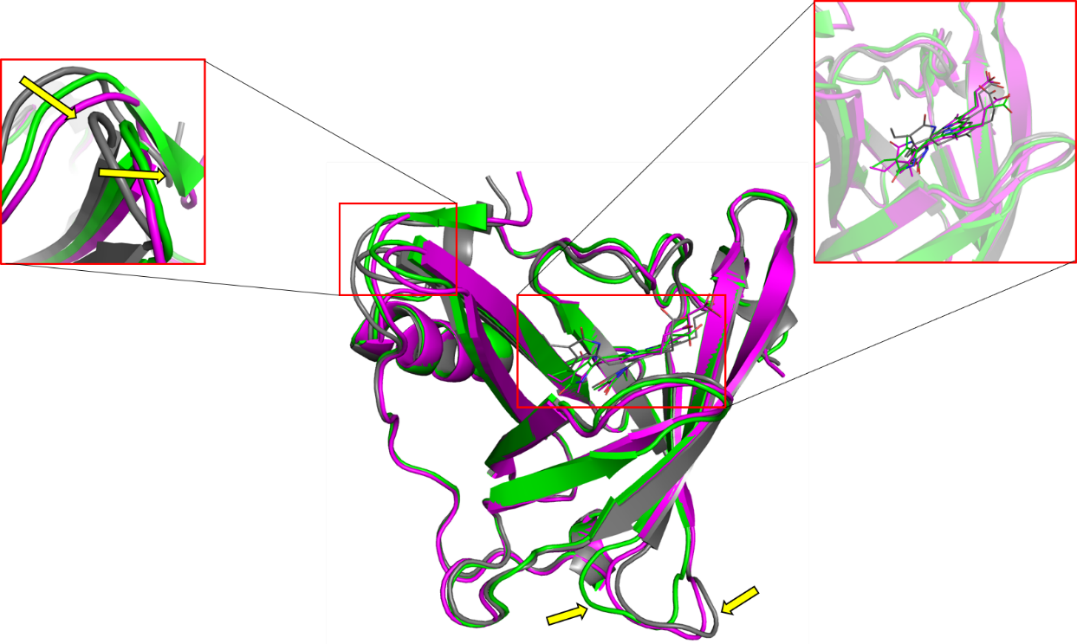

(C)

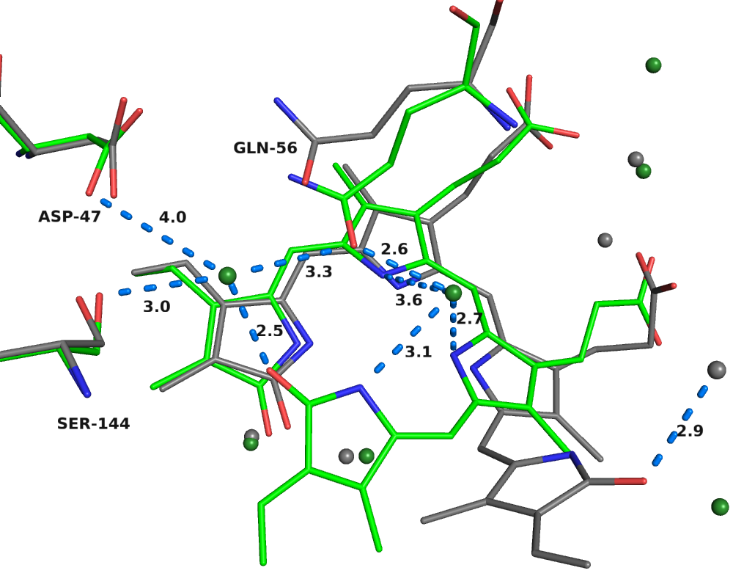

**Table 1. X- ray data collection and refinement statistics of monomeric and tetrameric variants of Sandercyanin**

| **Crystal** | **V71E -BLA complex (monomer)** | **L135E -BLA complex (monomer)** | **Y142A-BLA complex (tetramer)** |
| --- | --- | --- | --- |
| **PDB ID** | **7O2Y** | **7O3K** | **7O32** |
| **Data collection** | ID23-2, ESRF | BM14, ESRF | ID30A-1, ESRF |
| **Space group** | P 41 | P 41 | P 63 2 2 |
| **Unit cell dimensions** | 38.5 38.5 117.6 | 39.7 39.7 118.9 | 161.3 161.3 82.2 |
| **Resolution range (A^0^)** | 36.56 - 2.5 (2.59 - 2.5) | 39.65 - 2.75 (2.849 - 2.75) | 44.42 - 2.65 (2.745 - 2.65) |
| **Total reflections** | 41927 (3399) | 132365 | 363949 |
| **Unique reflections** | 6126 (685) | 4772 (488) | 18797 (1840) |
| **Multiplicity** | 6.8 | 27..0 | 19.0 |
| **Completeness (%)** | 99.70 (99.35) | 99.46 (96.63) | 99.18 (94.57) |
| **Mean I/sigma(I)** | 10.4 | 6.4 | 9.01 |
| **Wilson B-factor** | 36.55 | 43.69 | 27.30 |
| **R-factor** | 0.1732(0.1748) | 0.1971 (0.1964) | 0.2177 (0.4148) |
| **R-free** | 0.2535 (0.2415) | 0.2952 (0.3360) | 0.2809 (0.3663) |
| **Number of atoms:** |  |  |  |
| Macromolecules | 1281 | 1271 | 2580 |
| Ligands | 43 | 43 | 90 |
| Water | 51 | 40 | 90 |
| **Protein residues** | 166 | 166 | 338 |
| **RMS(bonds)** | 0.014 | 0.013 | 0.014 |
| **RMS(angles)** | 1.95 | 1.96 | 2.01 |
| **Ramachandran favored (%)** | 95.12 | 90.24 | 96.11 |
| **Clashscore** | 6.17 | 12.05 | 4.39 |
| **Average B-factor:** | 40.55 | 49.01 | 32.51 |
| Macromolecules | 40.63 | 48.93 | 38.64 |
| Solvent | 39.87 | 42.28 | 21.55 |

(Statistics for the highest-resolution shell are shown in parentheses)

**Figure S3. Spectral properties and structure of BLA- bound Y142A**

(A) Fluorescence and excitation spectrum of BLA- bound Y142A. Excitation spectra reveal similar spectral features as monomeric variants (V71E and L135E) shown in Figure 1C. (B) Structure of BLA- binding pocket of Y142A after refinement with partially occupied ZZZsss conformation show water molecules (86 and 93, dark-teal) overlapped with waters 67 and 68 (magenta) in V71E.

(A)

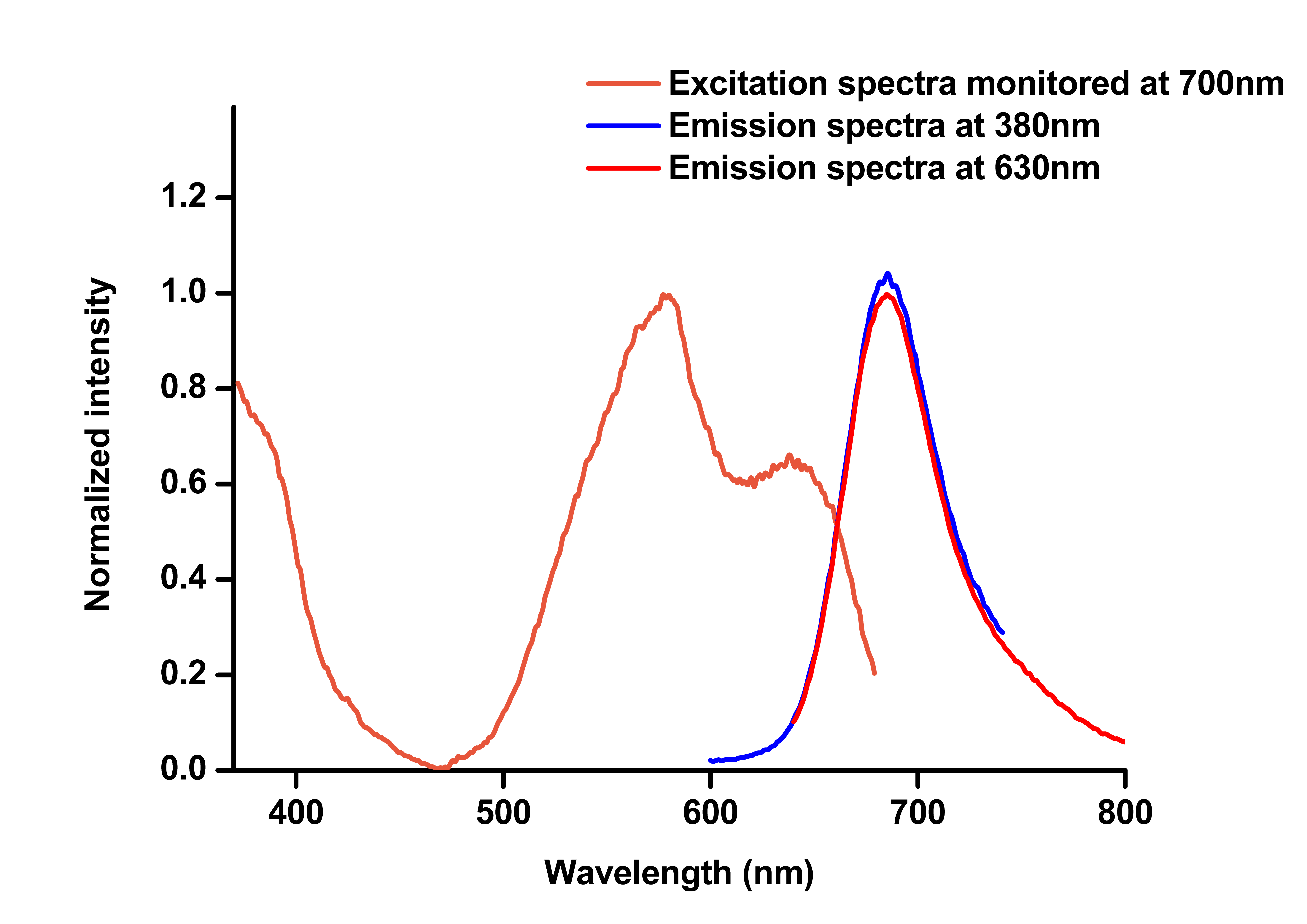

(B)

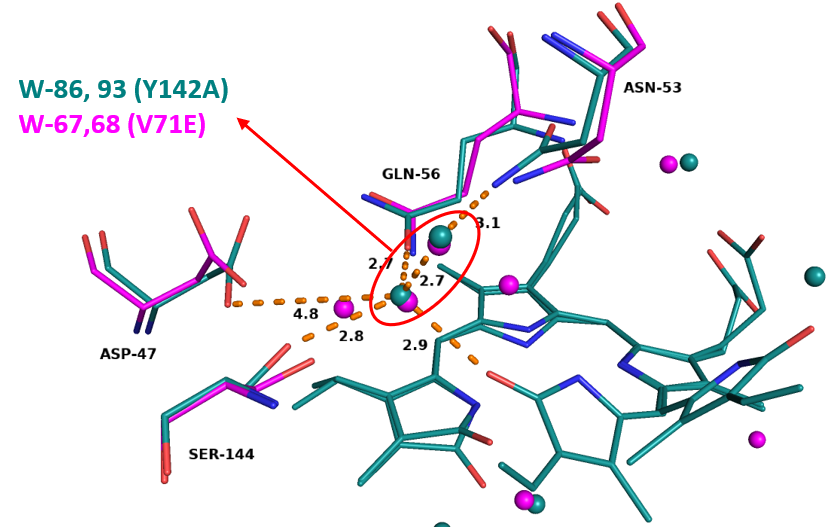

**Table S3.** Normal mode assignment of resonance Raman bands of free BLA, wild-type and Tyr-142-Ala (Y142A) Sandercyanin.

| BLA IXa | | | | | | Sandercyanin | | | | Assignment | Localization of normal mode |
| --- | --- | --- | --- | --- | --- | --- | --- | --- | --- | --- | --- |
| Experiment,  resonance Raman | | Computed ^a^ | | _b_ | _c_ | Wild type | | Y142A mutant | |  |  |
| λ_exc_ = 405 nm | λ_exc_ = 532 nm | Normal Raman | Raman scattering  activities | Raman, Soln.  1064 nm | SERS, Soln.,  514 nm | λ_exc_ = 405 nm | λ_exc_ = 532 nm | λ_exc_ = 405 nm | λ_exc_ = 532 nm |  |  |
| cm^-1^ | cm^-1^ | cm^-1^ | A^4^/AMU | cm^-1^ | cm^-1^ | cm^-1^ | cm^-1^ | cm^-1^ | cm^-1^ |  |  |
| 1699, w | 1696, vw | 1693 & 1703 | 870 & 920 | 1699 | - | 1695, w | 1695, m | 1699, w | 1693, vw | C=O str. | loc., A-ring, D-ring |
| - | - | 1655 | 12867 | - | - | - | - | - | - | C-C str. ring + C_4_=C_5_ str., | loc., A-ring |
| 1636, sh, m | 1634, sh, m | 1672 | 3695 | - | - | 1646, sh, w | 1647, sh | 1644, sh, m | 1648, w | C-C str. ring + C_15_=C_16_ str. + C-C str. CH-CH_2_ | loc., D-ring |
| 1616, vs | 1618, sh, s | 1644 | 70815 | - | 1604 | 1625, vs | 1625, vs | 1620, vs | 1625, m | C-C str. ring + C_15_=C_16_ str. | loc., D-ring |
| - | - | 1616 | 3055 | - | - | - | - | - | - | C-C str. ring + C_9_=C_10_ str. | loc., A-ring, B-ring |
| 1588, sh, s | 1591 | 1603 | 9283 | 1593 | - | 1596, sh, m | 1596, m | 1591, sh, m | 1597, vw | C-C str. ring + C_9_=C_10_-C_11_ str. | loc., A-ring, B-ring |
| - | - | 1595 | 5690 | - | - |  |  |  |  | C-C str. ring + C-C str. CH-CH_2_ | loc., D-ring |
| 1528, w | 1528, w | 1552 | 3801 | - | 1544 | 1538, w | 1538, m | 1537, vw | 1539, vw | in-plane NH be., ring breath. | deloc. |
| 1464, vw | 1467 | 1524 | 13522 | 1470 | 1468 | 1468, m | 1477, m | 1471, w | 1471, m | C-C str. aliph. + C-N str. ring + CH_3_ def. | loc., C-ring |
| 1432, w | 1434 | 1494/1499 | 635/17419 | - | - |  | 1440, m | 1439, vvvw | 1440, vvw |  | - |
| - | - | 1395 | 2529 | - | - | 1406, vvw | 1406, m | - | 1405, vw | CH3 def. + CH be. + ring breath. | loc., C-ring, D-ring |
| 1384, vw | 1386 | 1386 | 763 | 1384 | 1379 | - | 1387, m | 1386, vvw | 1388, vw | CH3 def. + CH be. + C-C str. ring. + NH be. | loc., A-ring |
| 1353, w | 1352 | 1371 | 5282 | - | 1352 | 1350, vw | 1355, m | 1353, m | 1356, vvw | CH3 def. + C-C str. ring + C-C str. aliph. | loc., A-ring, B-ring |
| 1322, w | 1321, m, sh | 1321 | 155 | - | - | - | 1327, sh, w | 1326, m | 1327, m | CH be. + CH2 wag. + NH be. | deloc. |
| 1301, vvw | 1302, vs | 1325 | 12340 | 1301 | 1300 | 1301, vw | 1303, s | 1302, vvw | 1302, m | NH be. + C-N str. ring + ring breath. | deloc. |
| - | - | 1298 | 3188 | - | - | - | 1286, s | - | 1285, m | NH be. + CH be. | deloc. |
| 1260, sh, m | 1262, vs | 1270 | 29828 | - | 1260 | 1263, m | 1263, s | 1265, m, sh | 1263, m | NH be. + ring breath. + CH be. | loc., C-ring, D-ring |
| 1239, m, sh | 1244, m, sh | 1259 | 5133 | 1248 | - | 1250, sh, vvw | - | 1248, s | 1244, vvw | NH be. + CH be. + CH2 wag. | deloc. |
| 1224, w, sh | 1226, m, sh | 1252 | 3085 | - | - | 1234, sh, vw | 1236, sh, vvw | 1232, sh, vw | - | NH be. + CH2 wag. | deloc |
| - | - | 1200 | 1458 | 1193 | 1161 | 1194, w | - | 1188, w | 1189, vvw | ring breath. + CH2 wag. + C-C str. aliph. | loc., D-ring |
| 1168, w | 1170, w | 1187 | 707 | - | - | - | 1168, w | 1174 | - | ring breath. + CH2 wag. | loc., C-ring |
| 1139, vvw | 1138, vvw | 1170 | 309 | - | - | - | 1141, m | - | 1143, vvw | ring breath. + CH be. | loc., A-ring |
| 1118, vw | 1120, vvw | 1134 | 977 | - | - | 1118, m | 1114, vw | 1118, m | 1117, vvw | NH be. + ring breath. + CH be. + CH3 def. | deloc. |
| - | 1094, w | 1107 | 331 | 1095 | 1085 | 1097, sh, vvw | 1099, m | 1098, vvw | 1099, m | C-C str. + C-N str. + NH be. +CH be. | loc., D-ring |
| - | - | 1077 | 676 | - | - | - | 1068, sh | 1065, vw | 1068, m | CH2 wag. + C-C str. aliph. + CH3 def. | - |
| 1055, vvw | 1052, w | 1043-1057 | - | - | 1042 | 1059, w | 1052, m | 1057, m | 1052, m | CH3 def. | - |
| - | - | 1012 | 122 | - | - | - | 1014, vw | - | - | CH3 def. | - |
| - | - | 1000 | 1454 | - | - | - | 1004, w | - | - | CH2 wag. + ring breath | loc., D-ring |
| 992, w | 996, vw, sh | 1007 | 1604 | - | 993 | 996, w ^$$^ | 994, vvw | 993, m | - | CH3 def. + ring breath. | loc., A-ring |
| 966, w | 972, m | 981 | 4124 | 972 | - | - | 968, m | 966, m | 968, vw | CH2 wag. + ring breath. + CH3 def. | loc., C-ring |
| - | - | 954 | 191 | - | - | - | 956, sh | - | 958, sh, vvw | CH2 wag. + ring breath. + CH3 def. | loc., B-ring |
| 923, vvw | 926, w | 931 | 661 | - | 936 | 931, w | 933, m | 935, vw | 933, vw | NH be. + ring breath. + CH be. + C_9_C_10_C_11_ be. + C_4_C_5_C_6_ be. | loc., B-ring |
| - | - | 901/904 |  | - | 909 | - | 919, vw | - | - | Ring def. + C_14_C_15_C_16_ be. | deloc. |
| - | - | 881 | 746 | 885 | - | - | 899, w | - | - | ring str. | loc., A-ring |
| not obs. | 873, vvw | 883 | 375 | - | - | 872, vvw | 871, w | 874, vvw | 872, w | CH wag. | - |
| - | - | 865 | 2767 | 835 | 842 | 857, vw | 853, w | - | 852, w | HOOP, C_15_-H | loc., C_15_-H |
| 825, w | 828, vw | 816 | 7 | - | - |  | 822, sh | 835, sh, vw |  | Out-of-plane ring def. | loc., D-ring |
| - | - | 846 | 86.5 | 818 | - | 811, m | 809, m | 820, m | 821, w | CH2 wag. + OCO be. | - |
| - | - | 792 | 196 | 790 | - | - | 796, sh, vw | - | - | ring breath. | deloc. |
| - | - | 787 | 137 | 783 |  | - | - | - | - | Out-of-plane ring def. | loc., A-ring |
| not obs. | 761, vvw, * | 777 | 159 | 767 | - | - | 769, vvw | - | 775, vvw | Out-of-plane ring def. + CH2 wag. + C-C str. | loc., A-ring |
| - | - | 718 | 1134 | 722 | - | - | 722, sh, vw | 717, sh, m | 720, sh, vw | Out-of-plane ring def. + CH2 wag. + ring str. | loc., B-ring, C-ring |
| 708, w, * | 710 | 711 | 249 | 717 | - | - | 711, sh, w | - | 708, sh, w | CH2 wag. + ring str | deloc. |
| - | - | 702 | 1281 | 709 | - | 705, m | 700, sh | - | - | out-of-plane NH be. + ring def. | deloc. |
| - | - | 697 | 1609 | 688 | - |  | 686, m | 688, m | 684, vw | in-plane ring def. | deloc. |
| 672, w, * | 677 | 683/687 | 412/1110 | 679 | 680 | 672, m | 673, sh, w | 672, sh, vw | - | out-of-plane ring def. | deloc. |
| 647, sh, vw | 650, sh, v  w | 634 | 28 | 655 | 642 | 654, sh, vw | 655, m | 653, vvw | 654, sh, w | In-plane ring def. | deloc. |
| not obs. | 610, vvw | 617 | 143 | 620 | - | 610, vvw | 612, w | 609, vw | 610, vw | In-plane ring def. + out-of-plane NH be. | - |
| - | - | 584 | 62 | - | - | - | 581, sh, vw | 588, vw | 590, vw | In-plane ring def. | deloc. |
| 544, vvvw |  | 543 | 221 | - | - | 550, m | - | 550, vvw | - | ring str. + C-C str. aliph. | loc., A-ring |
| - | 505, vvw | 512 | 57 | 511 | - | 510, vw | - | 508, m | 510, sh, w | ring def. + CH2 wag. | loc., B-ring + C-ring |
| - | - | 472 | 105 | - | 488 | 486, vw | 488, vw | 487, m | 489, m | out-of-plane ring. def. | loc., A-ring + B-ring |
| - | - | 464 | 276 | - | - | 463, vvw | 464, sh, vw | 458, m | - | out-of-plane NH be. | loc., D-ring |

^a^ Computed with RB3LYP/6-31+G(2d,p) on BLA anion; b ref (36) ; c ref (37); abbreviations; s, strong; vs, very strong; m, medium; sh, shoulder; w, weak; vw, very weak; vvw, very very weak; Aliph., aliphatic; str., stretching; be., bending; def., out-of-plane deformation; as., asymmetric; wag., wagging or out-of-plane bend; loc., localized on a ring; breath., in-plane-breathing; ^$$^ 992 cm-1 band corresponding to A-ring localized mode is used to normalize the 405 nm excited spectrum in Figure S5 panel A.

**Table S4.** TD computed lowest excited states^a^; excitation energy in nm, and oscillator strengths of wild-type and Y142A Sandercyanin. The corresponding experimental absorption maxima are mentioned in parenthesis.

| S_1_ | | | | | | |
| --- | --- | --- | --- | --- | --- | --- |
|  | wt | Y142A | | | | |
| Model | Structure | Clusters (from MD) | | | | |
|  | PDB: 5EZ2 | A | B | C | D | E |
| Population (%) | - | 5.3 | 14.6 | 12.3 | 7.4 | 61.8 |
| Excitation Energy (nm) | 568  (675) | 638.4  (720) | 577.8  (660) | 580.5  (660) | 577  (660) | 640.7  (720) |
| Oscillator Strength | 0.45 | 0.16 | 0.26 | 0.30 | 0.29 | 0.25 |

^a^ Excited states are computed with ONIOM (TD-CAMB3LYP:Amber) method and 6-31+G(d) basis set on high-QM layer, along with electronic embedding of MM changes. (See Methods for more details)

**Figure S4**. Vibrational assignment of RR spectrum of free BLA. Spectrum of BLA (a) computed on BLA anion at B3LYP//6-31+G(2d,p) with PCM implicit solvation; Experimental spectrum of BLA (b) with 532 nm excitation at pH 7; and (c) with 405 nm excitation at pH 7; (d) three normal modes of BLA anion with computed harmonic vibrational wavenumber, experimental wavenumber at 532nm excitation is mentioned in parenthesis. Details of the normal modes are described in Table S3 in Supplementary information.

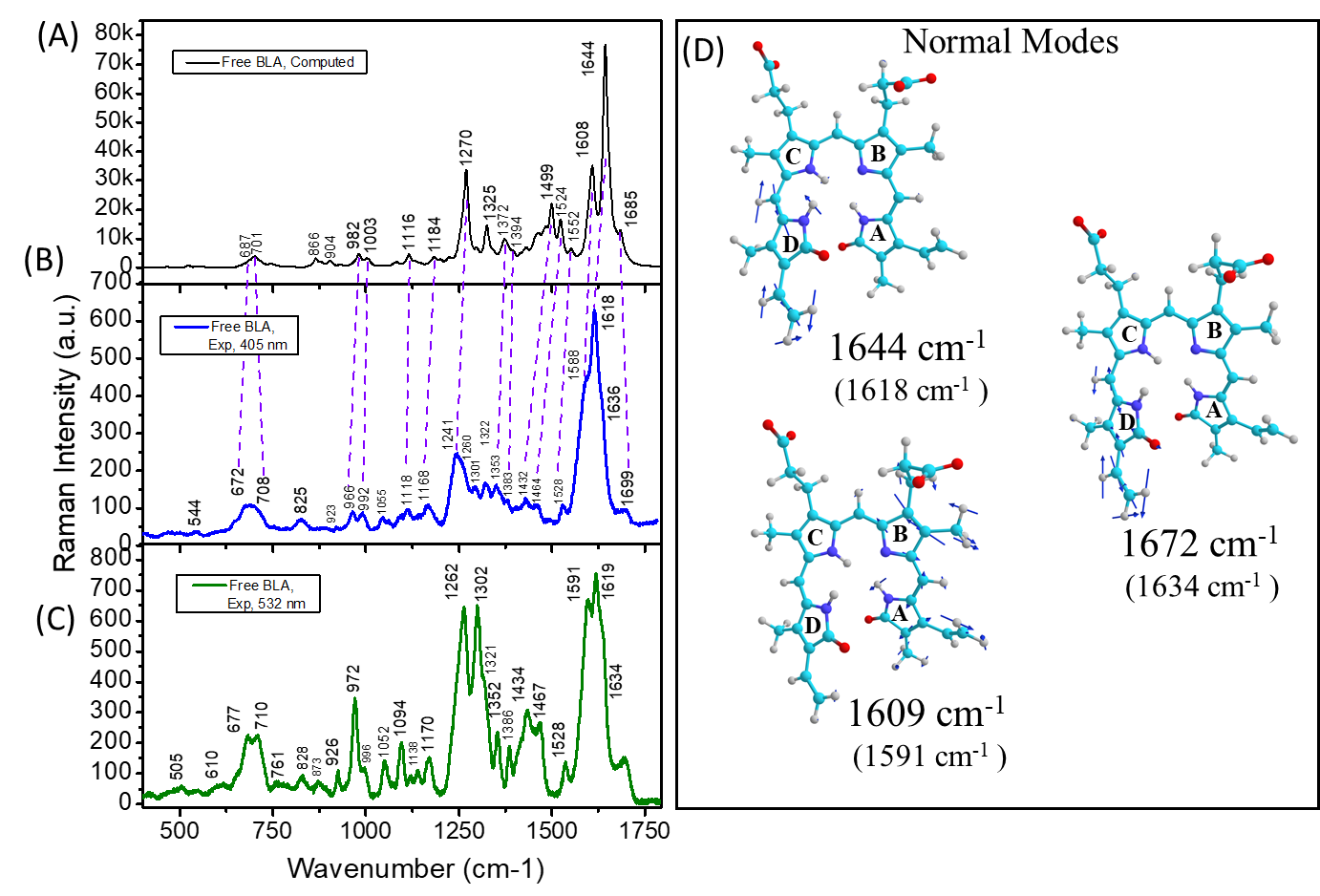

**Figure S5**. (A) Resonance Raman spectra of wild-type Sandercyanin (blue) and Y142A (orange) Sandercyanin at 405 nm excitation. Both spectra are normalized at 992 cm-1 (marked by arrow, and described in Table S3), a band exclusively originating from Ring-A vibration without ant contribution from Ring-D. (B) QM/MM computed normal Raman spectrum of wild-type Sandercyanin (green) and Y142A (brown) Sandercyanin. The computed spectra show in Y142A (brown), normal Raman activity of the BLA modes involving D-ring vibrations within 1550 – 1750 cm-1 decreases compared to that in wild-type protein (green). See SI Appendix for details of the QM/MM simulation. We note that, our QM/MM computation of Raman activity does not take into account resonance effect. Computation of RR spectra explicitly needs computation of normal modes displacement on potential energy surface of electronic excited state. While being done for small molecules, implementation of computation of RR intensity within QM/MM approach with more than 100 atoms in the QM region is extremely challenging.

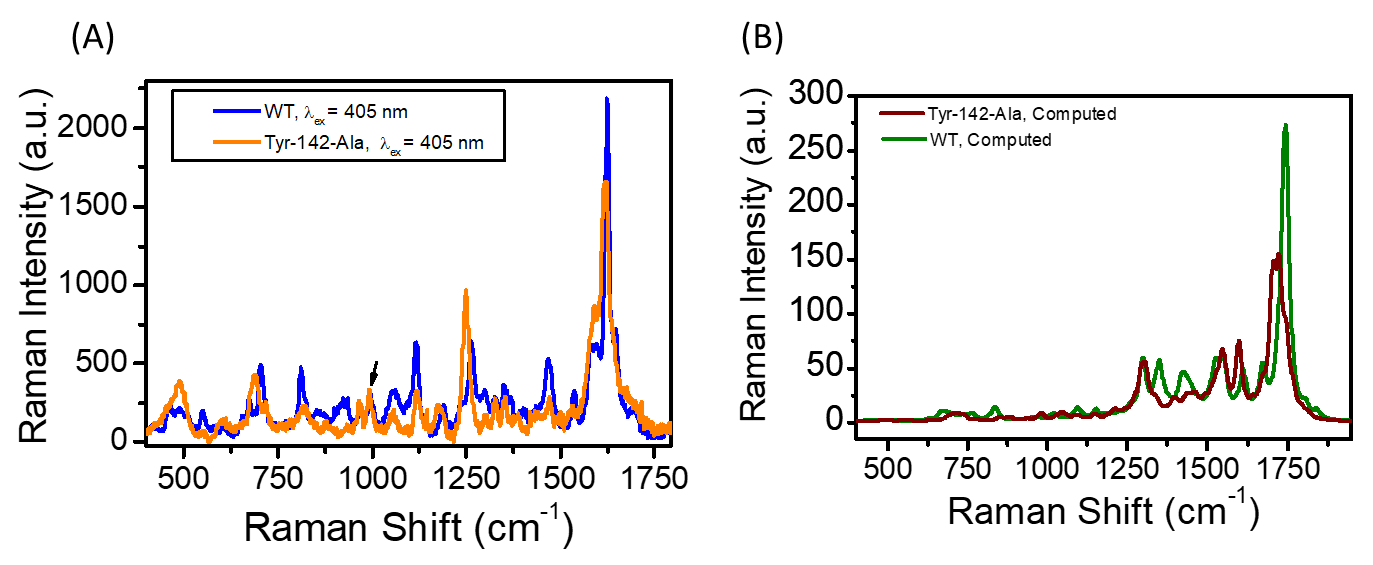

**Figure S6. Classical MD simulation detects most stable BLA conformation in Y142A Sandercyanin along with conformation of the active site residues.** (A-E) MD predicted BLA structures representing five clusters based on RMSD of all heavy atoms of BLA and residues within 4 Å. (F) Superposition of the all five structure depicts heterogeneity in the position of Phe-55 and His 108, and Tyr-116 stabilizing BLA in the pocket of Y142A Sandercyanin. (G) The important molecular interactions that stabilize the most populated ensemble of Y142A structure (shown in Panel E). TD-ONIOM(QM:MM) computed first excited state energies are described as λ_abs_, and the associated photo-states (SFr or Sr) are also mentioned for all five clusters in panel A-E. See Table S4 for excitation energies, and method section for details of computations.

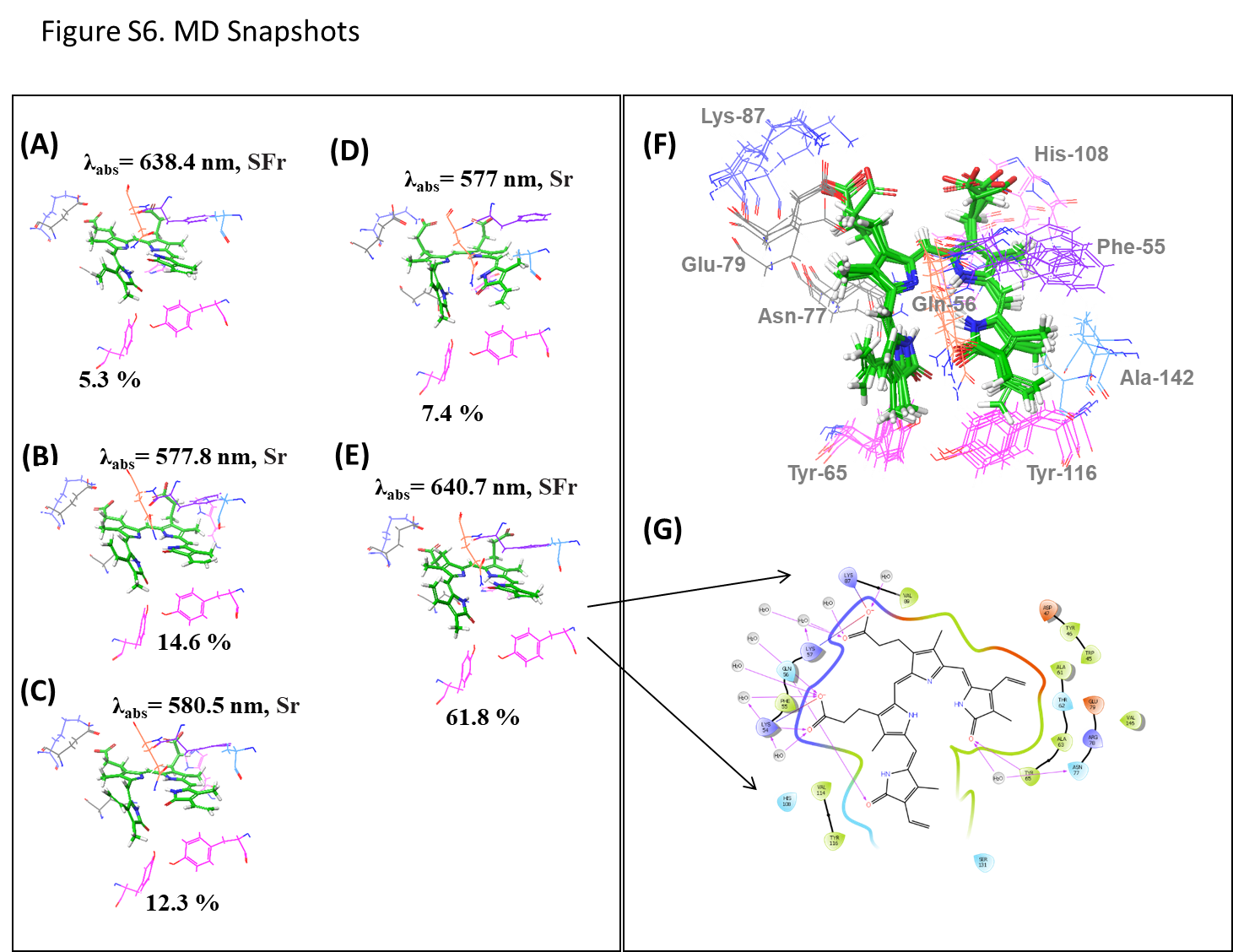

**Figure S7. Monte Carlo-Stochastic Dynamics (MC-SD) simulation probes most probable BLA conformation in Y142A Sandercyanin.** (A) MC/SD predicted BLA structures representing four clusters based on torsional RMSD around N-C14-C15=C16 dihedral (inset). The corresponding dihedral angle is mentioned on each representative structure in degree, and population is also mentioned in percentage; (B) Definition of the dihedral between C-ring and D-ring used in panel A; (C) Superimposed most populated conformation of BLA of Y142A from MC-SD simulation (in green), and that in crystal structure of V71E monomeric variant (in orange) of Sandercyanin. RMSD for heavy atoms of BLA are mentioned in case of superposed structures in panel C.

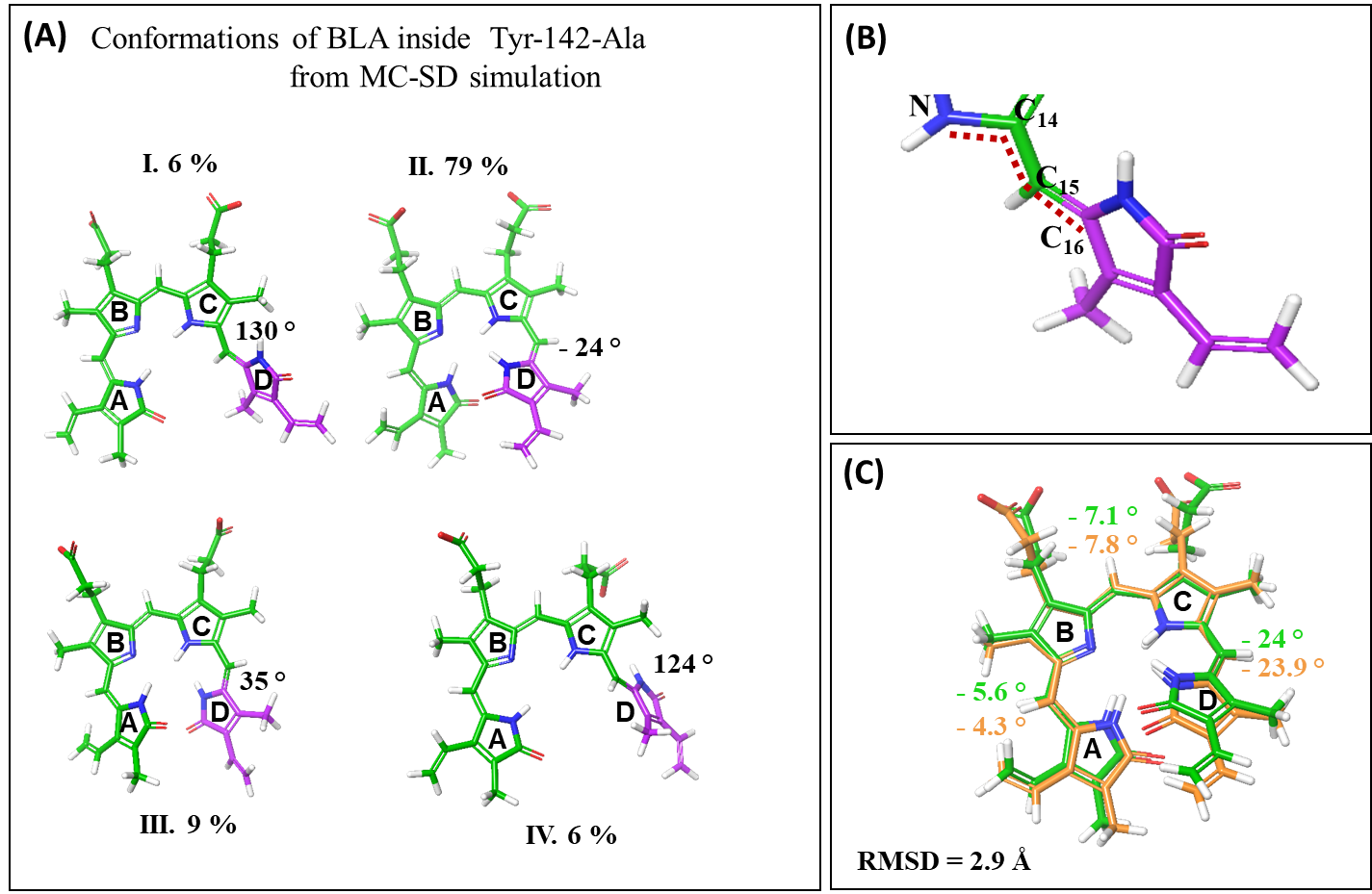

**Figure S8.** **Flow chart of theoretical methods applied in this study, starting from wild-type crystal structure of apo- Sandercyanin**. Inset shows definition of link atoms in two-layers ONIOM(QM:MM) calculations.

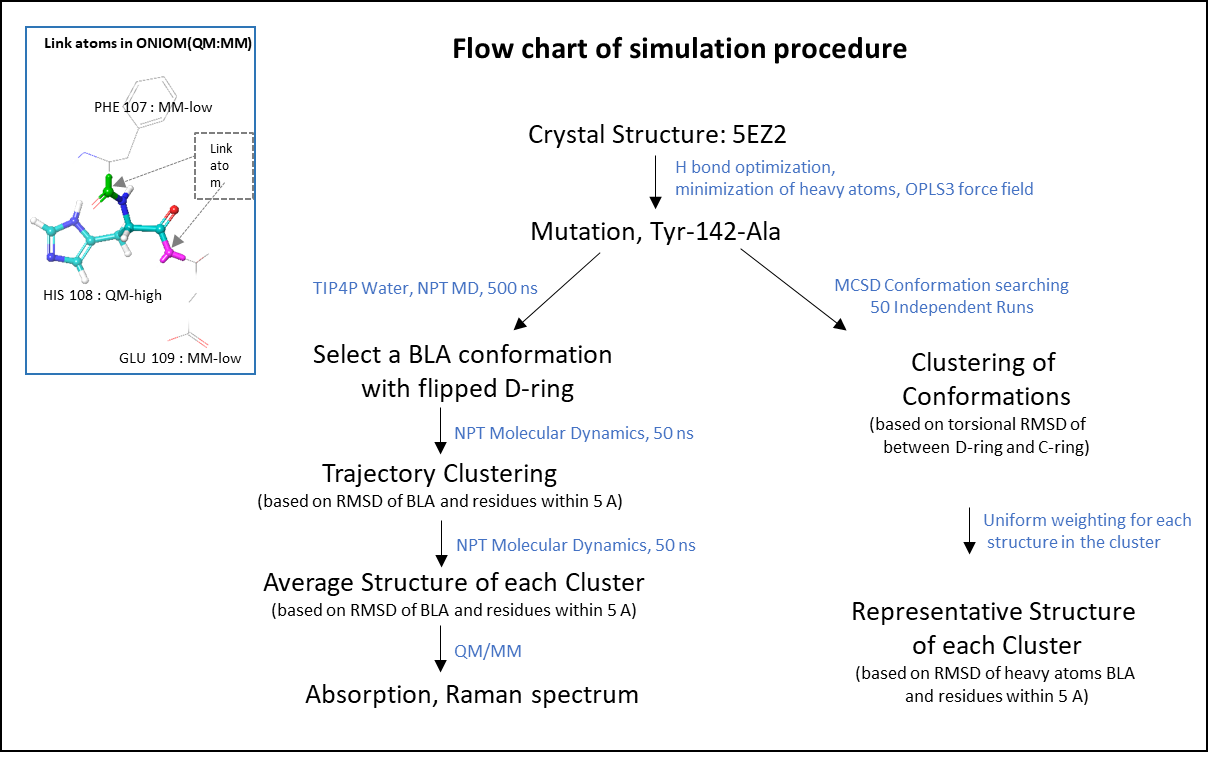
